# Development of larvae of the Australian blowfly, *Calliphora augur* (Diptera: Calliphoridae), at constant temperatures

**DOI:** 10.1101/2021.01.19.427229

**Authors:** Donnah M. Day, Nathan J. Butterworth, Anirudh Tagat, Gregory Markowsky, James F. Wallman

## Abstract

*Calliphora augur* (Diptera: Calliphoridae) is a common carrion-breeding blowfly of forensic, medical and agricultural importance in eastern Australia. Despite this, detailed information on the developmental biology of *C. augur* is lacking. Here, we present the first comprehensive study on the development of all three larval instars of *C. augur*, fed on sheep’s liver, at constant temperatures of 15, 20, 25, 30 and 35°C. We provide thermal summation models describing instar duration, as well as 95% prediction intervals for larval length at each constant temperature, enabling the age of larvae of *C. augur* to be estimated from their developmental stage and their average length. These data provide a basis for the application of this species to forensic investigations.

## Introduction

Forensic entomology is the study of insects as applied to legal issues. It may relate to medicolegal events, especially murders, for the purpose of uncovering information useful to an investigation ^1^. This is because decomposing vertebrate remains provide a temporary micro-habitat and food source that is attractive to a variety of insects and other invertebrates ^2^. Arthropods constitute a major element of this fauna, with insects, especially flies and their larvae, being the predominant group. Blowflies (Diptera: Calliphoridae) are ubiquitous and are typically the first insects to visit a dead body, often well before it is discovered by humans ^3^.

Data on the development of blowflies and other important carrion flies can help provide an estimate of the minimum time since death (or minimum post-mortem interval: mPMI), especially when combined with information on the succession patterns of insects on corpses^4^. Current approaches to age estimation are based on species-specific rates of development such as length, weight, and developmental duration in relation to temperature. While measures based on length and weight are valuable, developmental duration is the most widely used method of age estimation for two primary reasons; 1) because the duration of each stage of larval development (referred to as ‘instars’) is less variable than larval size (particularly in response to differences in diet)^5^; and 2) because instar stage is not affected by specimen shrinkage during preservation^6^. Once these measurements are obtained for the entire period of larval development, a variety of models can be used to estimate mPMI; including curvilinear regression ^8^, isomegalen and isomorphen diagrams ^9^, and thermal summation models ^10,11^. Of these, the most widely used are thermal summation models that incorporate temperature (usually as degree-hours) and time spent in each developmental stage ^12^.

*Calliphora augur* Fabricius is a common blowfly species found in eastern Australia ^13^, the larvae of which are among those most commonly collected from forensic cases, including from inside buildings ^14^. It is ovoviviparous ^15^, and also an agent of myiasis in sheep and other animals ^16–18^. Adult females prefer fresh carcasses ^19^, but will also larviposit on carcasses in active decomposition (i.e., 10-15 days old) ^20^. While previous studies have considered the larval development of *C. augur* ^16,20–24^ none have made fine-scale size measurements across a range of constant temperatures, or used thermal summation models to estimate mPMI. This lack of accurate developmental data means that at present, *C. augur* cannot be used reliably in a forensic context. To ameliorate this, we present the first comprehensive analysis of larval growth at constant temperatures of this important species. Our investigations are based on measurement of larval length throughout development, as well as observation of larval instar duration. Most importantly, we provide thermal summation models based on instar duration, and calculate prediction intervals based on larval length – for the first time allowing the application of this species to forensic investigations.

## Materials and methods

### Fly culture

A culture of *C. augur* was established from adults trapped at Wollongong, New South Wales (34° 25’ S, 150°53’ E). Experiments commenced using larvae of the F3 generation and were continued for 11 generations. This culture was refreshed regularly with wild-type (F0) individuals from the same source population as above and maintained with mixed ages of females and males. The average fecundity of each female was estimated by separating 20 females and transferring them to 50 mL specimen pots containing approximately 10 g of freshly defrosted sheep’s liver (defrosted to room temperature immediately prior to experiments). Larvae were counted the following day. Larvae and adults were kept in a temperature-controlled room at 25±3.5°C and ambient humidity, with a 12:12 light:dark regime incorporating a 15 minute ‘dusk’ and ‘dawn’ transition period of low light. Adult flies were maintained in square plastic cages measuring 330 mm long, 220 mm wide and 250 mm high (external dimensions). The flies in each cage were provided with water and sugar *ad libitum*, and chopped sheep’s liver in a small plastic weigh boat for larviposition as required. Larvae were reared in square white 2 L plastic containers, measuring 170 mm wide along each side and 90 mm deep. Most of the centre of the lid of each container was cut away and replaced with fine mesh to permit ventilation. In each container, larvae were sustained on sheep’s liver in the weigh boats placed atop a layer of wheaten chaff approximately 20 mm deep to enable pupariation. After pupariation, pupae were manually sorted from the chaff and transferred to a cage. Sheep’s liver was provided just prior to eclosion to provide females with a constant source of protein for ovary maturation ^20^.

### Development at constant temperatures

To assess rates of development at constant temperatures, freshly laid larvae were randomly collected from the main culture. Preliminary studies on larval density indicated that some endogenous heat generation occurred with as few as 25 larvae growing on 50 g of sheep’s liver. To ensure that the larvae developed at the ambient temperature being examined, each replicate (a plastic weigh boat containing larvae and liver defrosted to room temperature immediately prior to the experiment) was limited to 10 larvae. The growth period (between 0 and 600 hours) of each replicate of 10 larvae at a particular temperature was chosen by lottery. Between three and six replicates were used for each time interval (Supplementary Material 1). These replicates were grown at specific constant temperature regimes (15, 20, 25, 30 and 35°C ± 1.0 °C) and 60 ± 5% relative humidity, in an Axyos (Brisbane, Queensland, Australia) environmental cabinet. To ensure that the temperature inside the cabinet matched the set value, each replicate also contained a temperature logger (iButton; accuracy ±1.0 °C, resolution 0.5 °C; Maxim Integrated Products, Sunnyvale, CA, USA). Since only one cabinet was available, growth at each temperature was studied in turn. Larvae were left undisturbed in darkness in the environmental cabinet until collection, because disturbance can delay pupariation ^20,24^. To account for growth rates increasing with temperature, we sampled more frequently at higher temperatures. For 15°C, sampling was conducted every twenty-four hours, up to a total of 600 hours. For 20°C, sampling was conducted every six hours, up to a total of 216 hours. For 25, 30 and 35°C sampling was conducted every four hours, up to a total of 204 hours. The choice of these specific sampling frequencies was guided by the growth curves of O’Flynn^24,25^, and frequencies were kept relatively high to minimise error in the calculation of the prediction intervals.

### Sample collection, handling and preservation

At collection, all larvae were killed and fixed immediately by immersion in boiling water, dried with paper towel and preserved in 80% EtOH. Larvae were placed into glass vials posteriad to prevent head-curling ^26^ and length and instar were recorded.

### Larval length measurements

The body lengths of larvae were measured with the aid of a dissecting microscope and Mitutoyo Absolute digimatic digital callipers (Kawasaki, Japan) after the larvae had been in preservative for a minimum of ten days ^27^. Body length was measured as the distance, viewed laterally, between the most distal parts of the head and the last abdominal segment. The lengths of pupae were also measured with callipers. The dataset consists of the measurements of the lengths of larvae grown at temperatures of 15°C, 20°C, 25°C, 30°C, and 35°C. Since no larvae were measured more than once, the length data for the various time intervals should be regarded as independent.

### Larval instar durations

The instar of each larva was also recorded when it was measured as either 1st, 2nd, or 3rd, allowing for the calculation of thermal summation models based on instar duration. We estimated a thermal summation model separately for each stage of development. We estimated two parameters of the thermal summation model given by (DT = k + t D), where DT is the product of D, development duration (in hours) and T, environmental temperature in degrees celsius. The parameters, k (sum of effective temperatures) and t (lower developmental threshold) were estimated using ordinary least squares regression as well as reduced major axis regression (RMA) (Ikemoto and Takai, 2000) using the statistical software Stata 16.1. The coefficient on development duration is recovered as *t* and the constant is recovered as *k*.

## Results

On average, gravid females laid 58 ± 14 eggs. In total, 3,200 individual larvae (320 replicates of 10 larvae) were measured across a range of temperatures (15, 20, 25, 30 and 35°C ± 1.0°C) (Figure 1). Regarding the timing of developmental stages, the duration of each instar varied substantially between temperatures (Figure 2). As was expected, the average development time of each instar decreased with increasing temperature up to 30°C but increased slightly for third instars at 35°C. To determine the precise relationships between instar duration and degree-hours, thermal summation models were calculated using ordinary least squares regression (Figure 3) and reduced major axis regression (Figure 4). From these models, the sum of effective temperatures (*k*) and lower developmental threshold (*t*) were calculated for each instar for both the ordinary least squares regression (Table 1) and reduced major axis regression (Table 2).

**Table 1.** Summary of development constants for C. augur for three developmental stages, based on ordinary least squares regression. Sum of effective temperatures (*k*) and lower developmental threshold (*t*) shown as means with standard errors (coefficient of determination (R^2^) and degrees of freedom (Df) and p-values are provided).

| Stage | Temperature range | $R^2$ | Df | p-value | $k (\pm SE)$ | $t (\pm SE)$ |
| --- | --- | --- | --- | --- | --- | --- |
| 1 | 15–35 | 0.9045 | 526 | < 0.001 | $19.5135 \pm 2.5992$ | $19.202 \pm 0.2717$ |
| 2 | 15–35 | 0.7744 | 586 | < 0.001 | $361.332 \pm 10.889$ | $11.543 \pm 0.257$ |
| 3 | 15–35 | 0.7379 | 2068 | < 0.001 | $1072.928 \pm 27.335$ | $13.283 \pm 0.174$ |

**Table 2.** Summary of development constants for C. augur for three developmental stages based on the reduced major axis regression. Sum of effective temperatures (*k*) and lower developmental threshold (*t*) shown as means. Coefficient of determination (R^2^) and degrees of freedom (Df) and p-values are provided.

| Stage | Temperature range | $R^2$ | Df | p-value | $k$ | $t$ |
| --- | --- | --- | --- | --- | --- | --- |
| 1 | 15–35 | 0.951 | 526 | < 0.001 | 14.216 | 20.188 |
| 2 | 15–35 | 0.7744 | 586 | < 0.001 | 303.242 | 13.114 |
| 3 | 15–35 | 0.7379 | 2068 | < 0.001 | 792.078 | 15.462 |

**Figure 1.**
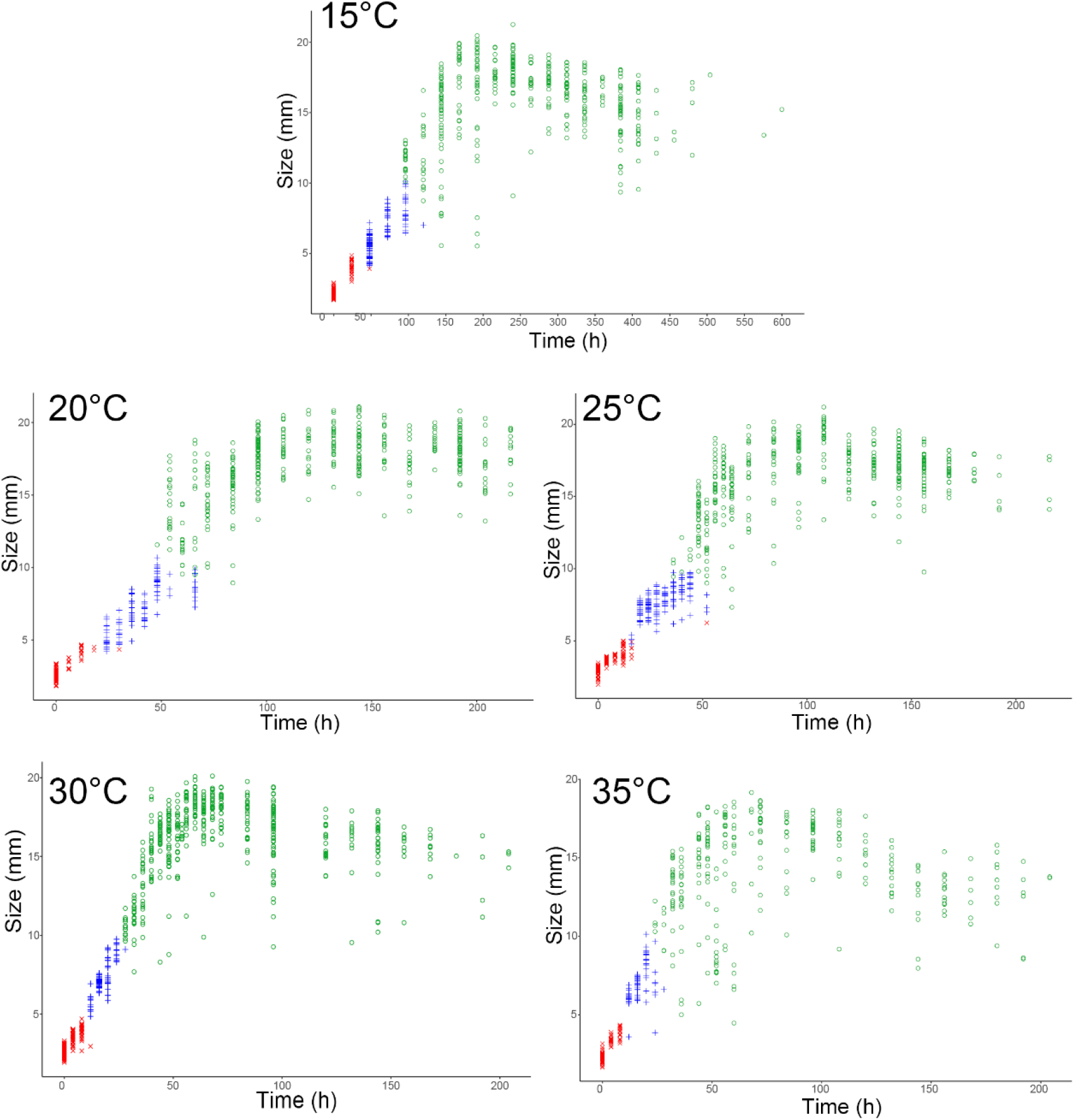
Growth curves of *Calliphora augur* larvae at constant temperatures. Data points represent individual larvae at each treatment time. First-instar larvae are represented by the ×, second-instar by the +, and third-instar by the o. The first plot (15°C) has a different scale on the x-axis.

**Figure 2.**
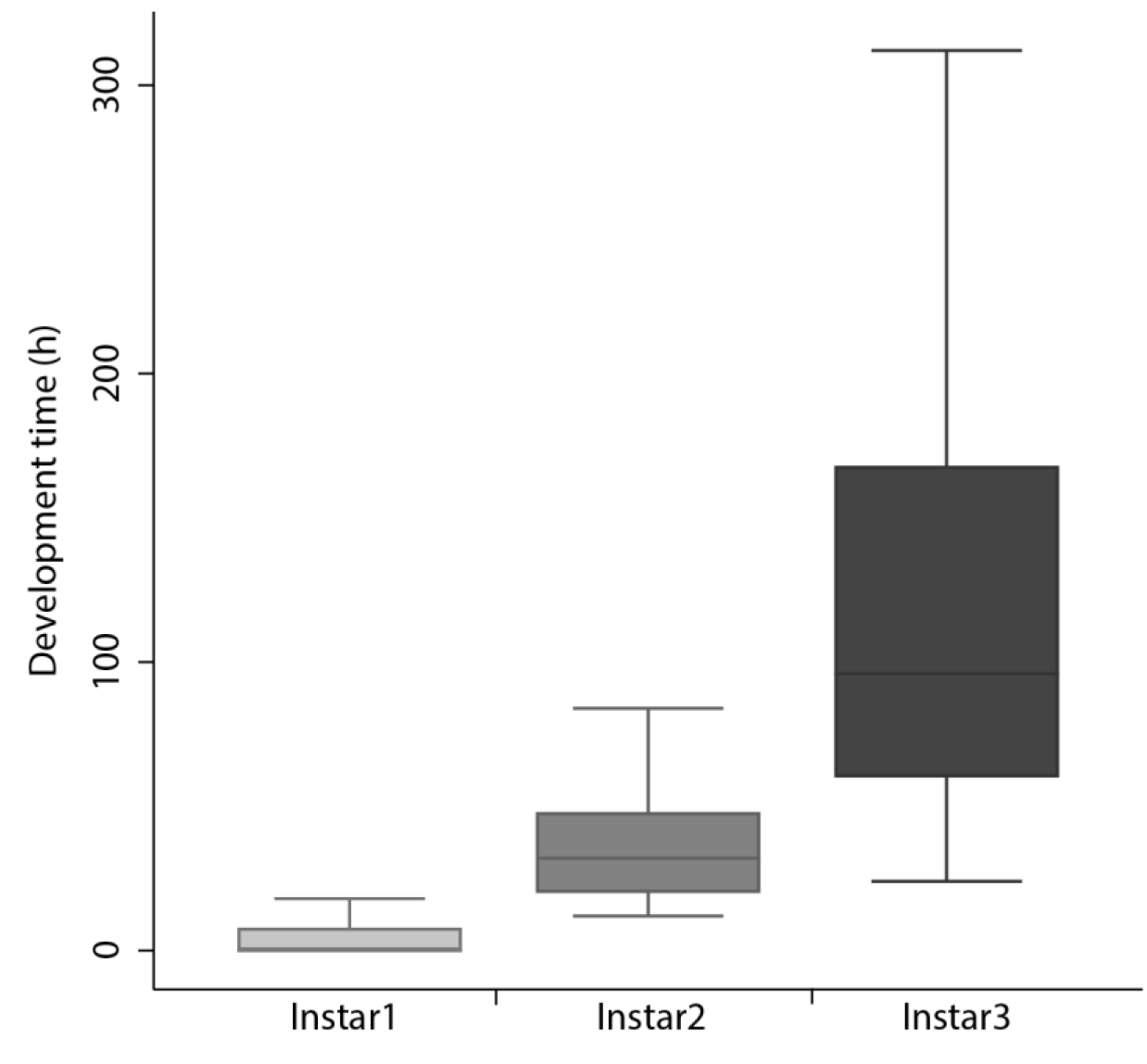
Observed range of instar durations of *Calliphora augur* larvae over five experimental treatments (15, 20, 25, 30, 35°C). The horizontal lines within boxes represent median values. The upper and lower boxes indicate 75^th^ and 25^th^ percentiles.

**Figure 3.**
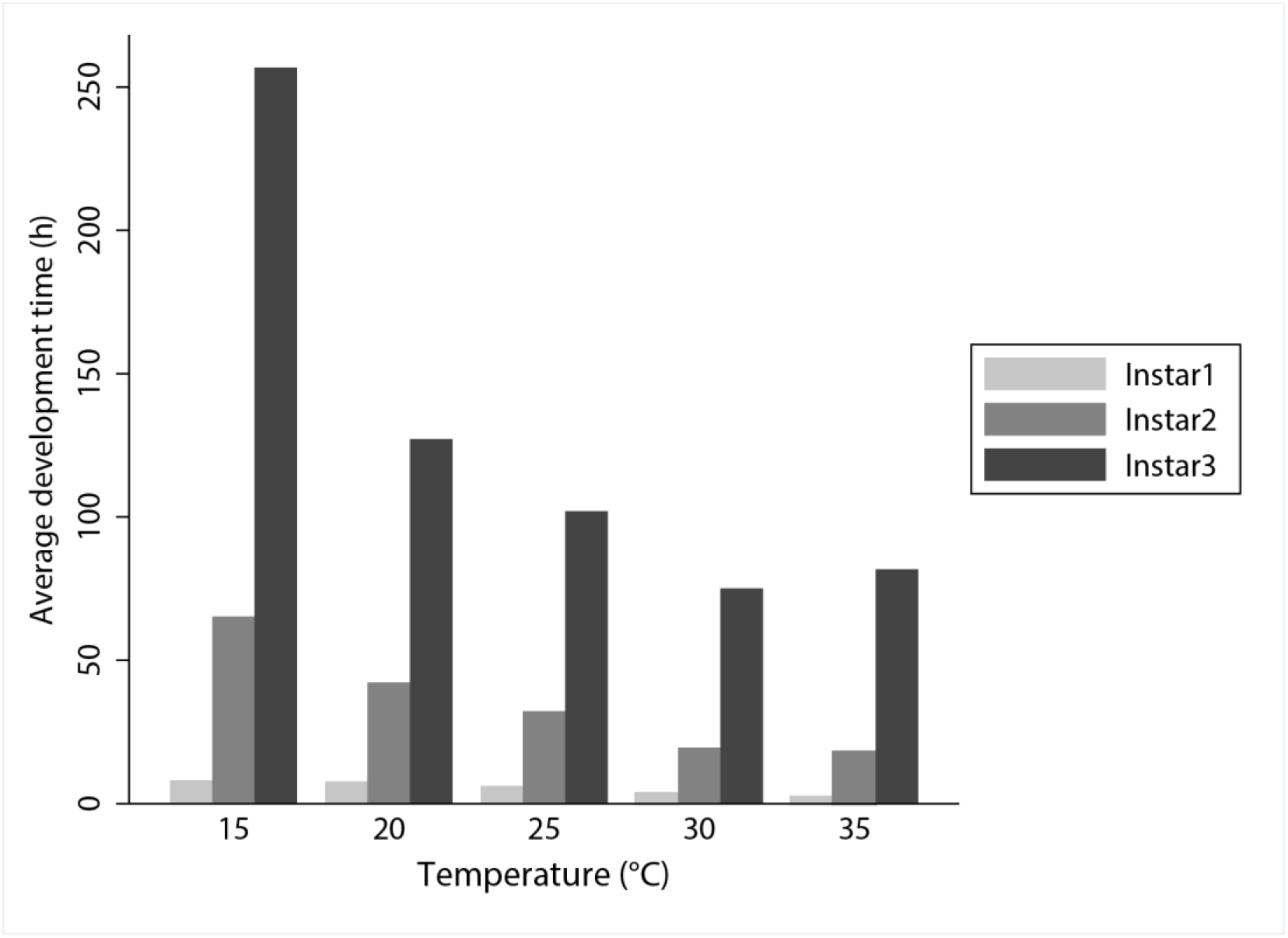
Bar plots of mean instar duration (hours) of all three instars of *Calliphora augur* across five experimental treatments (15, 20, 25, 30, 35°C).

**Figure 4.**
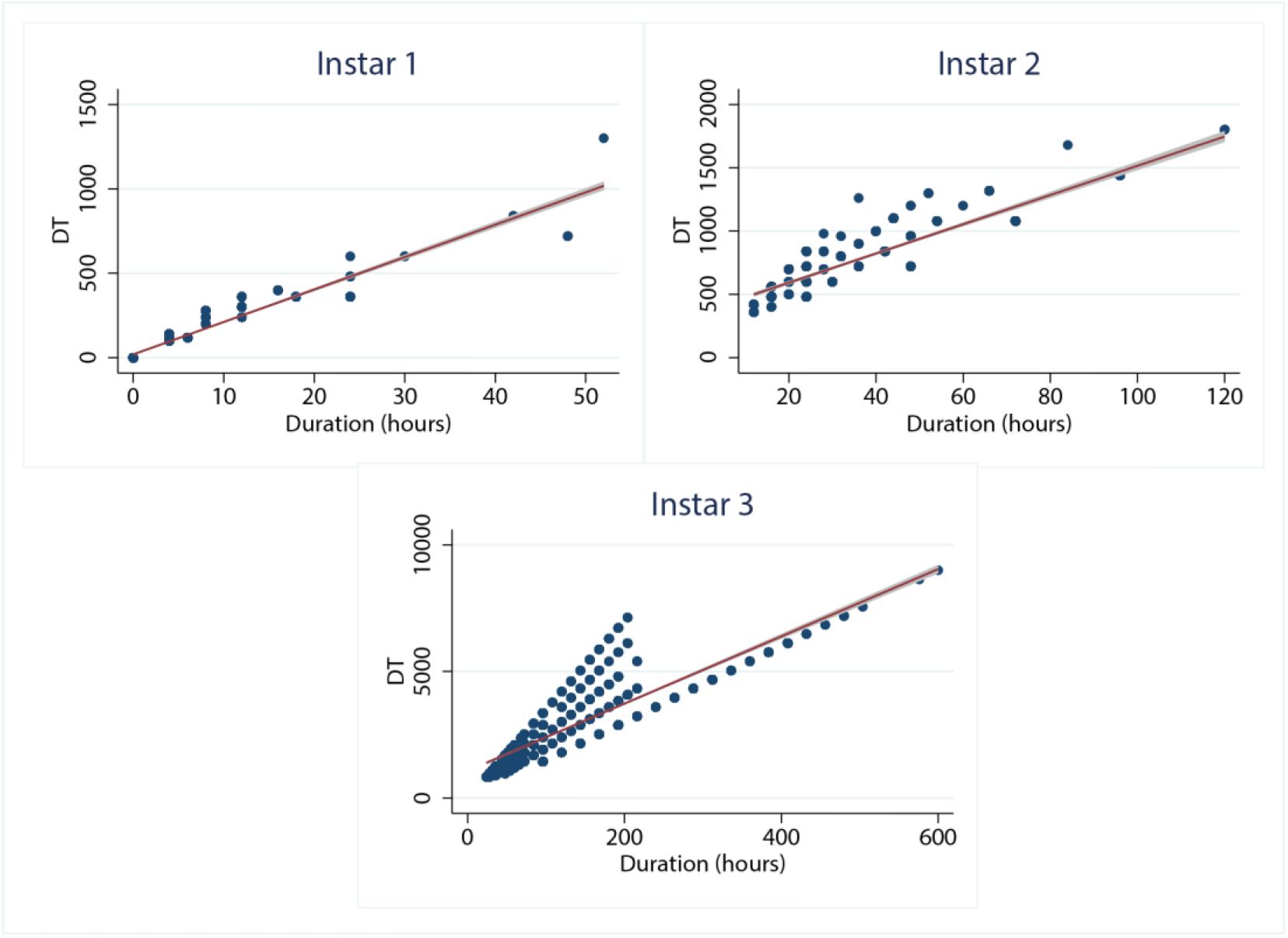
Ordinary least squares regression for the duration of development of each instar stage of *Calliphora augur*. DT represents the time in hours to reach the stage multiplied by the constant rearing temperature.

**Figure 5.**
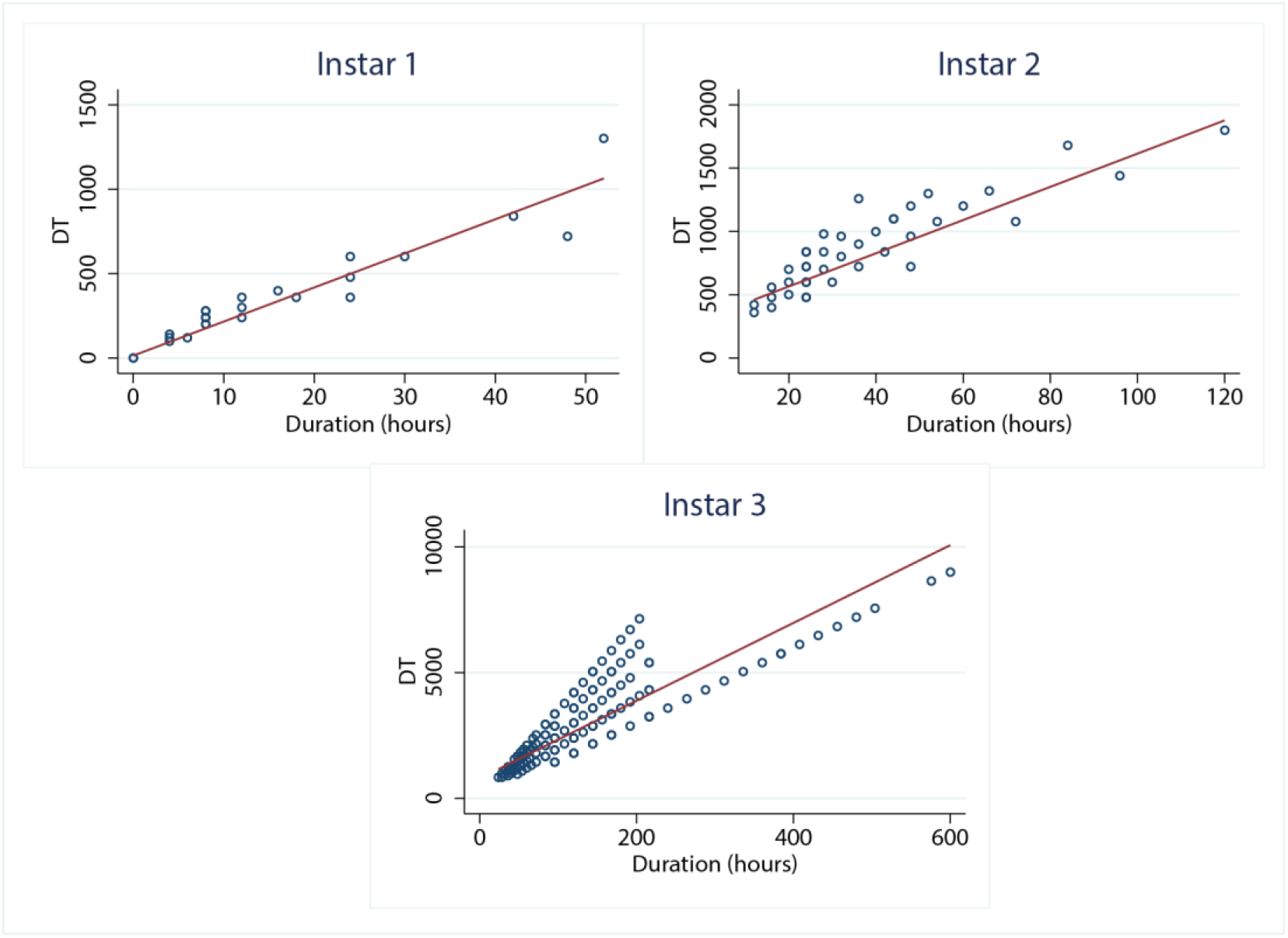
Reduced major axis regressions for the duration of development of each instar stage of *Calliphora augur*. DT represents the time in hours to reach the stage multiplied by the constant rearing temperature.

From our measurements of larval size, we also provide 95% prediction intervals that can assist in the estimation of age (Supplementary Material 2). Growth curves revealed that first-instar larvae were largest on average when grown at 25°C. Similarly, second instars were also largest when grown at 25°C, but they also displayed a much smaller variation at this temperature compared with the other temperatures examined. Third-instar larvae were largest on average when grown at 20°C, but were above the overall mean (for third-instars) when growing at 15, 20, 25 and 30°C. The only temperature at which the average length of third instars dropped below their overall mean was at 35°C. Post-feeding third instars were largest on average when grown at 20°C.

## Discussion

This study provides the first detailed dataset of the development of the Australian carrion-breeding blowfly *Calliphora augur*. The results we provide here reinforce the previous work done on this species but go substantially further by providing a level of statistical confidence in the application of developmental data in forensic investigations.

### Fecundity

Robust and highly active first-instar larvae of *C. augur* hatch promptly after deposition, giving this species an ecological advantage over oviparous species for exploiting small, quickly perishable carcasses ^17^. Johnston &Tiegs ^30^ reported that *C. augur* is capable of depositing eggs or larvae, with the eggs usually hatching within six hours. In the present study, we noted that *C. augur* will normally deposit living larvae but will also lay soft infertile eggs for some days before normal larviposition occurs – similar to the observations of Mackerras ^20^ and O’Flynn ^25^. The average number of larvae per female has previously been noted as 50 ^20^ or 58 larvae ^17^. Callinan ^31^ recorded a range of between 22.5 and 98.3 larvae per female in a controlled environment and a range of between 0.3 and 45.2 larvae per female in a field environment. The average number of larvae deposited in the present study was 58 ± 14 per gravid female, which is consistent with these previous data.

### Development at constant temperatures

The outcomes of some other developmental studies have been limited by a less thorough approach than taken here. For example, Byrd and Butler ^32^ noted that their sampling method selected for the fastest growing larvae, which often indicates a shorter period than could otherwise be deduced from normal collection techniques at a crime scene. The influence of size extremes upon variance has also been considered by Wells &LaMotte ^7^. In the present study, we included larvae of all sizes to demonstrate the full range that may be encountered in crime scene samples. Variation in larval body length appeared to decrease steadily from 15 to 30°C and then increased again dramatically at 35°C. The decrease in variation with increasing temperature may relate to a more even growth rate at higher temperatures. However, the subsequent increase in variation at 35°C likely reflects the fact that the larvae are reaching their thermal limit, at which point some larvae may begin to suffer from increased heat and oxidative stress, hindering their growth and increasing the perceived variation in size^33^. The variation in overlap of the growth stages, particularly the transitional forms between instars, emphasises the importance of assessing crime scene samples for instar and/or transitional forms, and collecting an adequate number of specimens. While there has been comment on the value of transitional forms ^34^, for most blowfly species information on transitional forms is lacking.

The reader will note an interesting feature of the plots related to larval length, namely that the measured lengths increase linearly with time to a certain point, but then begin to decrease after that. This late-stage reduction in larval length is a result of post-feeding shrinkage that calliphorid larvae undergo after leaving the carcass, prior to pupariation. This post-feeding shrinkage can cause some difficulty in interpreting the age of larvae, because the smaller length of post-feeding larvae makes them appear younger. However, this can be solved if investigators record where larvae are collected, as post-feeding larvae will depart the carcass and move into the surrounding substrate. If larvae are collected away from the carcass, it can be presumed that they are in the post-feeding stage and as such, post-feeding confidence intervals should be used to infer their age. Furthermore, post-feeding larvae will rapidly evacuate the crop prior to pupariation, and thus crop dissection can be performed to check for the presence of food particulates. If no food particulates are found, then it can be assumed that the larvae are post-feeding. For a more thorough discussion of this technique see Archer et al. ^35^.

While some other previous studies have included data on the development of *C. augur* ^16,20–24^, they were usually conducted as parts of larger projects and provide insufficient detail for inferring an mPMI. For example, Fuller ^16^ only reported the average length of the maggot when full grown (18 mm), with which our findings concur. Mackerras ^20^ referred only to complete development, with 21-22 d recorded for growth in summer conditions in an insectary and 18-20 d for development in a room at 20°C. Levot et al. ^23^ examined weights of *C. augur* larvae at 27-28°C and found the time to reach maximum larval weight to be 65.5 hours. While direct comparisons cannot be drawn with this, we observed that the time to reach maximum larval length at 30°C was 72 h. The techniques of Levot et al. ^23^ were very different to ours, with a large mass of larvae feeding on excess liver and each sampling event removing some individuals. It is possible that the larvae in their experiment were actually exposed to higher temperatures than the ambient of 27-28°C via massing, and that the feeding mass was better able to liquefy the substrate, and therefore grow more quickly, than the larvae in the current study. Comprehensive work by O’Flynn ^24,25^ examined larval growth at temperatures between 5 and 45°C but did not produce sufficient data for explicit mPMI predictions. At 5°C, the minimum period from larviposition to commencement of wandering was 54 d; some larvae survived for 110 d but none pupated.

### Larval temperature preference

It is well known that blowfly larvae show species-specific temperature preferences, which are behavioural adaptations that maximise developmental rate, growth quality, and survival ^36^. Examination of instar size and duration across temperatures in our study indicates that the larvae of *C. augur* also experience optimal growth at specific temperatures related to their growth stage. In terms of size, the only temperature at which the average length of third instars dropped below their overall mean was at 35°C, and post-feeding third instars were largest on average when grown at 20°C. Regarding instar duration, the shortest developmental times were observed at 35°C for first and second instars, and at 30°C for third instars – but the most substantial reduction in development time for all instars was seen between 15 and 20°C. In addition, it appears that the lower developmental threshold (*t*) for *C. augur* lies between 11 and 20°C (depending on instar). Altogether, this suggests that the development of *C. augur* is likely optimal around 30°C, but suffers at temperatures above this. Importantly, the species may be capable of completing development at temperatures as low as 11°C and can comfortably complete development at 20°C – which is in line with this species remaining active during the Australian winter.

### Environmental conditions and larval growth

Importantly, it is well established that the growth and survival of blowfly larvae is dependent upon their food source. For example, studies on *Calliphora vicina* and *Calliphora vomitoria* have shown substantial differences in larval growth between mixed minced meats and beef liver ^37^ and there are differences in larval growth between livestock and human tissue^38^. While the measurements we provide here are based on *C. augur* fed on sheep’s liver, these data are assumed to apply to the equivalent growth in human bodies – an assumption that holds true for comparisons between *C. vicina* larvae fed on porcine and human tissue^5^. Nevertheless, this assumption remains to be tested with *C. augur*, and age calculations based on data from animal tissue may lead to inaccurate mPMI estimates ^37,38^. For most forensically important species, including *C. augur*, there remains a pressing need for broad-scale studies that compare growth rates between a range of tissues from different animal taxa^37,5^.

In addition to this, it is important to note that the larvae reared in our developmental experiments were kept in darkness without a day/night light cycle. It is well known that photoperiod can influence larval growth rate and development^39,40^, and as such, our data may not reflect the developmental rate that would occur under a 12:12 day night cycle. Nevertheless, very few studies have explored the effect of complete darkness on insect larval development, so it is unclear to what extent this would influence developmental rate. In addition, many forensic cases concerning entomology involve corpses found in secluded places such as abandoned buildings, houses, and basements^41^ which are often exposed to very low levels of light (and in certain cases complete darkness). Future studies of *C. augur* development, and indeed blowflies more generally, will benefit from investigating how larval development differs under a range of lighting conditions (including full darkness, artificial light, and natural light).

## Conclusions

Overall, we provide prediction intervals based on larval length and thermal summation models based on instar duration, at a range of constant temperatures. This comprehensive dataset forms a basis for estimating the age of larvae of *C. augur*, for the first time enabling the application of this species to forensic investigations.

## Acknowledgements

We thank the University of Wollongong for financial assistance with this study.

## Disclosure statement

The authors have no conflict of interest to declare.

## Data availability statement

All data has been made available as supplementary material.

## Notes

### Competing Interest Statement

The authors have declared no competing interest.

