## Supplementary Material 2 for "Development of larvae of the Australian blowfly, *Calliphora augur* (Diptera: Calliphoridae), at constant temperatures"

Table 1. The 95% prediction intervals based on the developmental curves of *Calliphora augur* larvae at five different constant temperatures (15, 20, 25, 30, 35°C). *n* = sample number, $\bar{x}$*_exp_ =* sample mean, *s*^2^ = sample variance. *m = 5* interval is the prediction interval to be used for sample sizes between 5 and 29 larvae, *m = 30* interval is the prediction interval to be used for sample sizes larger than 30 larvae.

| **Time (hours)** | ***N*** | $\bar{x}$***_exp_*** | $\boldsymbol{s}^{\boldsymbol{2}}$ | ***m = 5* interval** | ***m = 30* interval** |
| --- | --- | --- | --- | --- | --- |
| **Instar 1, Temp 15 ^o^C** | | | | | |
| 0 | 59 | 2.1725 | 0.072 | (1.989,2.356) | (2.052,2.293) |
| 24 | 25 | 4.0148 | 0.240 | (3.636,4.393) | (3.741,4.289) |
| **Instar 2, Temp 15 ^o^C** | | | | | |
| 48 | 65 | 5.4798 | 0.530 | (4.986,5.974) | (5.159,5.801) |
| 72 | 26 | 7.3438 | 0.782 | (6.666,8.022) | (6.856,7.832) |
| 96 | 21 | 8.2752 | 1.286 | (7.366,9.184) | (7.602,8.948) |
| **Instar 3, Temp 15 ^o^C** | | | | | |
| 96 | 21 | 11.7024 | 0.632 | (11.065,12.340) | (11.231,12.174) |
| 120 | 17 | 11.9724 | 4.935 | (10.096,13.849) | (10.543,13.402) |
| 144 | 51 | 14.3841 | 10.767 | (12.105,16.663) | (12.868,15.901) |
| 168 | 25 | 17.3384 | 5.108 | (15.593,19.084) | (16.075,18.602) |
| 192 | 42 | 16.7124 | 13.344 | (14.117,19.308) | (14.949,18.476) |
| 216 | 19 | 17.7626 | 1.108 | (16.899,18.627) | (17.114,18.411) |
| 240 | 43 | 17.9709 | 3.062 | (16.731,19.211) | (17.131,18.811) |
| 264 | 20 | 17.149 | 2.339 | (15.909,18.389) | (16.225,18.073) |
| 288 | 31 | 17.0787 | 1.668 | (16.119,18.038) | (16.403,17.754) |
| 312 | 32 | 16.5694 | 1.934 | (15.542,17.597) | (15.849,17.290) |
| 336 | 29 | 16.0203 | 2.029 | (14.950,17.090) | (15.260,16.780) |
| 360 | 8 | 16.5700 | 0.607 | (15.696,17.444) | (15.837,17.303) |
| 384 | 41 | 14.7044 | 4.616 | (13.173,16.236) | (13.661,15.748) |
| 408 | 25 | 14.8204 | 4.069 | (13.263,16.378) | (13.693,15.948) |
| **Instar 1, Temp 20 ^o^C** | | | | | |
| 0 | 45 | 2.5538 | 0.164 | (2.268,2.839) | (2.361,2.746) |
| 6 | 11 | 3.3409 | 0.104 | (3.027,3.655) | (3.088,3.594) |
| 12 | 13 | 4.2562 | 0.135 | (3.919,4.593) | (3.990,4.522) |
| **Instar 2, Temp 20 ^o^C** | | | | | |
| 24 | 18 | 5.1522 | 0.519 | (4.553,5.752) | (4.699,5.605) |
| 30 | 13 | 5.5415 | 0.538 | (4.869,6.214) | (5.011,6.072) |
| 36 | 20 | 7.0295 | 0.865 | (6.276,7.783) | (6.468,7.591) |
| 42 | 19 | 7.1032 | 0.404 | (6.581,7.625) | (6.712,7.495) |
| 48 | 22 | 9.0773 | 0.772 | (8.380,9.774) | (8.564,9.590) |
| 66 | 10 | 8.5380 | 0.643 | (7.727,9.349) | (7.876,9.200) |
| **Instar 3, Temp 20 ^o^C** | | | | | |
| 54 | 18 | 14.4444 | 3.65 | (12.855,16.034) | (13.243,15.646) |
| 60 | 15 | 11.6647 | 1.906 | (10.456,12.874) | (10.728,12.601) |
| 66 | 14 | 14.8379 | 10.474 | (11.943,17.733) | (12.575,17.101) |
| 72 | 27 | 14.6637 | 3.818 | (13.177,16.151) | (13.598,15.729) |
| 84 | 38 | 15.6639 | 3.861 | (14.249,17.079) | (14.692,16.636) |
| 96 | 64 | 17.7798 | 2.149 | (16.784,18.776) | (17.132,18.428) |
| 108 | 24 | 18.245 | 1.689 | (17.233,19.257) | (17.509,18.981) |
| 120 | 18 | 18.1456 | 2.534 | (16.821,19.470) | (17.144,19.147) |
| 132 | 27 | 18.6196 | 1.475 | (17.695,19.544) | (17.957,19.282) |
| 144 | 44 | 18.2527 | 2.302 | (17.181,19.325) | (17.528,18.977) |
| 156 | 19 | 18.3826 | 2.562 | (17.069,19.696) | (17.397,19.369) |
| 168 | 18 | 17.2128 | 2.393 | (15.926,18.500) | (16.240,18.186) |
| 180 | 16 | 18.6719 | 0.627 | (17.992,19.352) | (18.149,19.194) |
| 192 | 54 | 18.1091 | 1.673 | (17.216,19.002) | (17.518,18.700) |
| 204 | 16 | 16.8313 | 3.503 | (15.223,18.439) | (15.596,18.066) |
| 216 | 14 | 18.0257 | 2.066 | (16.740,19.311) | (17.021,19.031) |
| **Instar 1, Temp 25 ^o^C** | | | | | |
| 0 | 48 | 2.9033 | 0.121 | (2.660,3.147) | (2.740,3.066) |
| 4 | 27 | 3.6081 | 0.037 | (3.462,3.754) | (3.503,3.713) |
| 8 | 11 | 3.81 | 0.048 | (3.597,4.023) | (3.638,3.982) |
| 12 | 30 | 4.268 | 0.214 | (3.923,4.613) | (4.024,4.512) |
| 16 | 6 | 4.2917 | 0.178 | (3.732,4.852) | (3.807,4.777) |
| **Instar 2, Temp 25 ^o^C** | | | | | |
| 20 | 23 | 7.143 | 0.291 | (6.719,7.567) | (6.833,7.453) |
| 24 | 28 | 7.31 | 0.352 | (6.862,7.758) | (6.990,7.630) |
| 28 | 21 | 7.5662 | 0.864 | (6.821,8.311) | (7.015,8.118) |
| 32 | 18 | 7.9783 | 0.203 | (7.603,8.353) | (7.695,8.262) |
| 36 | 16 | 8.275 | 0.712 | (7.550,9.000) | (7.718,8.832) |
| 40 | 16 | 8.6106 | 0.564 | (7.965,9.256) | (8.115,9.106) |
| 44 | 17 | 8.7971 | 0.609 | (8.138,9.456) | (8.295,9.299) |
| **Instar 3, Temp 25 ^o^C** | | | | | |
| 48 | 34 | 13.6715 | 2.144 | (12.600,14.743) | (12.925,14.418) |
| 52 | 19 | 12.1405 | 2.881 | (10.747,13.534) | (11.095,13.186) |
| 56 | 29 | 15.8128 | 2.342 | (14.663,16.962) | (14.996,16.629) |
| 60 | 27 | 16.0007 | 4.955 | (14.307,17.695) | (14.787,17.214) |
| 64 | 27 | 14.7056 | 5.339 | (12.947,16.464) | (13.446,15.966) |
| 72 | 21 | 16.5886 | 5.495 | (14.710,18.467) | (15.197,17.980) |
| 84 | 24 | 17.3142 | 5.99 | (15.409,19.220) | (15.928,18.701) |
| 96 | 39 | 17.7841 | 2.745 | (16.595,18.973) | (16.970,18.599) |
| 108 | 24 | 18.8704 | 3.099 | (17.500,20.241) | (17.873,19.868) |
| 120 | 27 | 17.2167 | 1.342 | (16.335,18.098) | (16.585,17.848) |
| 132 | 28 | 17.1954 | 1.806 | (16.180,18.211) | (16.471,17.920) |
| 144 | 45 | 17.0431 | 2.249 | (15.986,18.100) | (16.331,17.755) |
| 156 | 33 | 16.5845 | 2.917 | (15.329,17.840) | (15.707,17.462) |
| 168 | 21 | 16.7257 | 0.818 | (16.001,17.451) | (16.189,17.262) |
| 180 | 7 | 16.8843 | 0.592 | (15.956,17.812) | (16.094,17.675) |
| 192 | 6 | 15.6000 | 2.257 | (13.606,17.594) | (13.873,17.327) |
| **Instar 1, Temp 30 ^o^C** | | | | | |
| 0 | 60 | 2.5583 | 0.104 | (2.338,2.779) | (2.414,2.703) |
| 4 | 28 | 3.5518 | 0.153 | (3.256,3.847) | (3.341,3.763) |
| 8 | 29 | 3.9228 | 0.168 | (3.615,4.231) | (3.704,4.141) |
| **Instar 2, Temp 30 ^o^C** | | | | | |
| 12 | 14 | 5.6279 | 0.278 | (5.156,6.100) | (5.259,5.997) |
| 16 | 29 | 7.0290 | 0.106 | (6.784,7.274) | (6.855,7.203) |
| 20 | 21 | 7.6262 | 0.946 | (6.847,8.406) | (7.049,8.203) |
| 24 | 13 | 9.0131 | 0.288 | (8.521,9.505) | (8.625,9.401) |
| **Instar 3, Temp 30 ^o^C** | | | | | |
| 28 | 13 | 10.5392 | 0.349 | (9.998,11.081) | (10.112,10.967) |
| 32 | 23 | 11.2465 | 1.399 | (10.317,12.176) | (10.567,11.926) |
| 36 | 25 | 13.1956 | 2.810 | (11.901,14.490) | (12.259,14.132) |
| 40 | 25 | 15.5992 | 2.760 | (14.316,16.882) | (14.671,16.528) |
| 44 | 30 | 16.1283 | 2.987 | (14.838,17.419) | (15.216,17.041) |
| 48 | 38 | 16.0895 | 3.748 | (14.695,17.484) | (15.131,17.048) |
| 52 | 27 | 16.4419 | 2.032 | (15.357,17.527) | (15.665,17.219) |
| 54 | 6 | 16.4567 | 0.141 | (15.958,16.955) | (16.025,16.888) |
| 56 | 25 | 17.7292 | 2.825 | (16.431,19.027) | (16.790,18.669) |
| 60 | 28 | 18.4364 | 0.625 | (17.839,19.034) | (18.010,18.863) |
| 64 | 34 | 17.4650 | 2.535 | (16.300,18.630) | (16.654,18.276) |
| 68 | 28 | 18.0279 | 1.991 | (16.961,19.094) | (17.267,18.789) |
| 72 | 23 | 18.0283 | 0.908 | (17.280,18.777) | (17.481,18.576) |
| 84 | 28 | 17.4275 | 1.600 | (16.471,18.384) | (16.746,18.109) |
| 96 | 53 | 16.3130 | 4.688 | (14.815,17.811) | (15.320,17.306) |
| 120 | 22 | 15.7395 | 1.246 | (14.854,16.625) | (15.088,16.391) |
| 132 | 12 | 15.7417 | 4.745 | (13.689,17.795) | (14.104,17.379) |
| 144 | 28 | 15.6121 | 3.940 | (14.112,17.112) | (14.542,16.682) |
| 156 | 10 | 15.3620 | 2.889 | (13.642,17.082) | (13.958,16.766) |
| 168 | 9 | 15.5289 | 0.800 | (14.581,16.477) | (14.745,16.313) |
| **Instar 1, Temp 35 ^o^C** | | | | | |
| 0 | 63 | 2.2735 | 0.082 | (2.079,2.468) | (2.147,2.400) |
| 4 | 16 | 3.3800 | 0.074 | (3.146,3.614) | (3.201,3.559) |
| 8 | 19 | 3.8411 | 0.129 | (3.546,4.136) | (3.62,4.062) |
| **Instar 2, Temp 35 ^o^C** | | | | | |
| 12 | 18 | 6.0122 | 0.462 | (5.447,6.578) | (5.585,6.440) |
| 16 | 27 | 6.8485 | 0.261 | (6.460,7.237) | (6.570,7.127) |
| 20 | 21 | 8.3519 | 0.920 | (7.583,9.121) | (7.783,8.921) |
| 24 | 8 | 6.8000 | 2.611 | (4.988,8.612) | (5.280,8.320) |
| **Instar 3, Temp 35 ^o^C** | | | | | |
| 28 | 5 | 10.4360 | 1.655 | (8.480,12.392) | (8.711,12.161) |
| 32 | 25 | 13.0576 | 3.477 | (11.618,14.498) | (12.015,14.100) |
| 36 | 20 | 11.9110 | 9.849 | (9.367,14.455) | (10.015,13.807) |
| 44 | 19 | 14.5695 | 8.228 | (12.215,16.924) | (12.803,16.336) |
| 48 | 27 | 15.2704 | 5.556 | (13.477,17.064) | (13.985,16.556) |
| 52 | 15 | 10.0413 | 9.612 | (7.327,12.756) | (7.939,12.144) |
| 56 | 20 | 16.1190 | 6.689 | (14.022,18.216) | (14.556,17.682) |
| 60 | 17 | 13.4012 | 22.225 | (9.418,17.384) | (10.367,16.435) |
| 68 | 6 | 15.8800 | 11.792 | (11.322,20.438) | (11.932,19.828) |
| 72 | 22 | 16.7327 | 3.414 | (15.267,18.198) | (15.654,17.811) |
| 84 | 14 | 15.7243 | 5.512 | (13.624,17.824) | (14.083,17.366) |
| 96 | 28 | 16.5311 | 0.826 | (15.844,17.218) | (16.041,17.021) |
| 108 | 14 | 15.7743 | 5.267 | (13.721,17.827) | (14.170,17.379) |
| 120 | 10 | 15.2280 | 1.652 | (13.928,16.528) | (14.166,16.290) |
| 132 | 13 | 13.9485 | 1.780 | (12.726,15.171) | (12.983,14.914) |
| 144 | 11 | 12.2182 | 5.073 | (10.025,14.411) | (10.449,13.987) |
| 156 | 13 | 12.9208 | 1.246 | (11.898,13.944) | (12.113,13.728) |
| 168 | 6 | 12.6067 | 2.463 | (10.523,14.69) | (10.803,14.411) |
| 180 | 9 | 13.4256 | 3.840 | (11.349,15.502) | (11.708,15.143) |
| 192 | 7 | 11.9214 | 5.784 | (9.021,14.821) | (9.451,14.392) |

Table 2. The sample means and variances of the growth of *Calliphora augur* larvae for cases which included between one and five measurements. Dots represent variances that could not be calculated due to there being only one sample.

| **Instar** | **Temp (^o^C)** | **Time** | ***n*** | $\bar{x}$***_exp_*** | ***s^2^*** |
| --- | --- | --- | --- | --- | --- |
| **1** | **15** | 48 | 1 | 3.910 | . |
| **2** | - | 120 | 1 | 7.020 | . |
| **3** | - | 432 | 4 | 14.203 | 3.890 |
| - | - | 456 | 2 | 13.340 | 0.168 |
| - | - | 480 | 4 | 15.385 | 5.512 |
| - | - | 504 | 1 | 17.680 | . |
| - | - | 576 | 1 | 13.400 | . |
| - | - | 600 | 1 | 15.230 | . |
| **1** | **20** | 18 | 2 | 4.395 | 0.026 |
| - | - | 30 | 1 | 4.350 | . |
| **2** | - | 54 | 3 | 8.707 | 0.584 |
| **3** | - | 48 | 1 | 11.570 | . |
| **1** | **25** | 52 | 1 | 6.250 | . |
| **2** | - | 16 | 3 | 5.093 | 0.099 |
| - | - | 52 | 3 | 7.490 | 0.381 |
| **3** | - | 36 | 3 | 10.150 | 0.373 |
| - | - | 40 | 2 | 10.955 | 2.856 |
| - | - | 44 | 3 | 10.940 | 0.125 |
| - | - | 216 | 4 | 16.050 | 3.518 |
| **1** | **30** | 12 | 1 | 2.960 | . |
| **2** | - | 28 | 1 | 9.100 | . |
| **3** | - | 180 | 1 | 15.030 | . |
| - | - | 192 | 4 | 13.663 | 5.704 |
| - | - | 204 | 4 | 15.003 | 0.241 |
| **2** | **35** | 28 | 1 | 6.61 0 | . |
| **3** | - | 24 | 4 | 11.023 | 0.723 |
| - | - | 204 | 2 | 13.760 | 0.003 |

**Utilizing the dataset**

Our dataset can provide potentially valuable information to an investigator by simply comparing the ratios of the various instars found at a crime scene with those obtained in our experiment.

For instance, supposing that a batch of larvae is recovered from a body from a location where the average temperature has been 15^o^C, and these larvae are roughly evenly split between instars 2 and 3, then consulting Figure 1 would give a range of the mPMI between 72 and 120 hours, with 96 hours being the best estimate. On the other hand, if only larvae in instar 1 are recovered from a body at an average temperature of 35^o^C, then the mPMI is confidently less than 12 hours, and so forth. This should help to establish the mPMI in certain circumstances, in particular when the percentages of the various instars are specific to a relatively small time interval, but is clearly not good enough in other instances, most notably those corresponding to longer mPMIs in which one is likely to find only larvae in instar 3. These cases require a more refined analysis, which we now provide.

It is assumed that an investigator is able to procure at least five larvae from a crime scene and is able to correctly record their average length, as we refer to in the ‘measurement and statistics’ section. We will provide 95% prediction intervals for the average length of the larvae for each combination of time, instar, and temperature; this is analogous to but different than giving confidence intervals, since we are predicting the mean value of a sample rather than the population mean. A detailed explanation of the process can be found in Geisser and Johnson ^28^, but for the benefit of the reader we briefly review the theory behind the method. Those unfamiliar with some of the standard distributions and facts from statistics theory used below are advised to consult almost any appropriate statistics textbook or online resource.

Suppose that, for a given time, instar, and temperature, the length of a maggot is normally distributed with mean *µ* and variance $\sigma^{2}$. We regard our experimental data as a sample from this population of size *N*, and wish to test the hypothesis that a sample procured from a crime scene by an investigator was taken from the same population. The crime scene sample is regarded as independent from the experimental sample, and let *m* be the number of larvae taken from the time scene. Let us label the sample mean from our experiment as $\bar{x}$_exp_ and from the crime scene as $\bar{x}$_cri_.

As is well known, $\bar{x}$_exp_ and $\bar{x}$_cri_ will be normally distributed random variables, each with mean *µ* but with variances σ^2^/*N* and σ^2^/*m* respectively. Since $\bar{x}$_exp_ and $\bar{x}$_cri_ are independent random variables, their difference $\bar{x}$_exp_ - $\bar{x}$_cri_ will also be normally distributed, but now with mean 0 and variance σ^2^/*N* + σ^2^/*m*.

It follows that $Z=(\bar{x}$_exp_ - $\bar{x}$_cri_)/$\sqrt{\sigma^{2}(\frac{1}{N}+\frac{1}{m})}$ is a standard normal (mean 0, variance 1). If we knew the population variance *σ*^2^, we could use a standard normal table to create 95% prediction intervals, but since we do not (as is typical) we will use the sample variance *s*^2^ from our experiment in its place. Now, $\frac{\left( N-1 \right)s^{2}}{\sigma^{2}}$ will have a χ^2^(*N*-1) distribution, and will be independent of *Z* (it is independent of $\bar{x}$_cri_ by assumption, and of $\bar{x}$_exp_ by the standard independence of sample mean and variance for normal random variables). Thus, $\frac{Z}{\sqrt{s^{2}/\sigma^{2}}}$ will follow a *t_n-1_* distribution. Replacing *Z* by the expression that defined it and cancelling the σ^2^ in the numerator and denominator, we see that $(\bar{x}$_exp_ - $\bar{x}$_cri_)/$\sqrt{s^{2}(\frac{1}{N}+\frac{1}{m})} \sim$ *t_n-1_*. Our 95% prediction interval for $\bar{x}$_cri_ is therefore $\bar{x}$_exp_ ± *t*_.025,N-1_ $\sqrt{s^{2}(\frac{1}{N}+\frac{1}{m})}$.

A few comments on the normality assumption made above are in order. It is commonly accepted that measurements such as the size of organisms are generally modelled well by normal distributions; indeed, we ran the standard Kolmogorov-Smirnov and Shapiro-Wilk normality tests on the data, and found that most of the data subsets for each time, instar, and temperature are consistent with an underlying normal distribution, with a small number of exceptions. Furthermore, even in the case of the exceptions, the *t*-test is known to be highly robust against non-normality, for the reason that even when the population does not follow a normal distribution the sample means $\bar{x}$_exp_ and $\bar{x}$_cri_ will still be approximately normally distributed ^29^. This assures us that the method described above is general and not dependent upon the normality assumption.

We present the prediction intervals in table form (Table 1), with a separate section for each combination of temperature and instar. Note that we have not provided tables for the intermediate stages (first- to second-instar, or second- to third-instar) since they do not appear in sufficient numbers to provide valuable prediction intervals; we have also removed from Table 1 the times at which there were fewer than five larvae of a particular instar present, for the same reason. The prediction intervals are given in the form (*a*,*b*), which is an equivalent notation to *a* ≤ $\bar{x}$_cri_ ≤ *b*. An investigator should make use of these tables by calculating the mean length of their sample of larvae, and then comparing this with the prediction intervals given in the table corresponding to the correct instar and average temperature; if the mean of the sample lies within the prediction interval for a given time, then that time should be regarded as a possible age for the larvae. We have provided prediction intervals for crime scene sample sizes *m = 5* and *m = 30*; the *m* = 5 interval should be used for sample sizes between 5 and 29, and the *m = 30* interval for any *m* over 30.

The investigator may also be interested in forming prediction intervals for time, temperature, and instar combinations that do not appear above. The table below (Table 2) may be of some use if the combination appears there, but there are still some combinations (for example, temperature 35^o^, instar 3, time 76 hours) for which we have no data. As a simple rule of thumb, in this case, if the nearest times for which there are data (here 72 and 84 hours) include $\bar{x}$_cri_ then the time in question should be regarded as possible. If the investigator requires a precise prediction interval then we recommend interpolating the means and variances linearly from the nearest points in time to estimate the mean and variance at the point in question, and taking the degrees of freedom to be equal to the minimum of the degrees of freedom at the nearest points (this is a slight simplification of the method used in Wells & Lamotte, 1995). For example, in our case of temperature 35^o^C, instar 3, time 76 hours, we look at the closest time points for the given temperature and instar, which are at 72 and 84 hours. The means are at 16.733 and 15.724, respectively, and the variances are 3.414 and 5.512. We therefore estimate the mean and variance at 76 hours to be $\left( \frac{2}{3} \right)16.733+\left( \frac{1}{3} \right)15.724=16.397$and $\left( \frac{2}{3} \right)3.414+\left( \frac{1}{3} \right)5.512=4.113$; note that the values at 72 hours are weighted double those at 84 hours, because 76 is twice as far from 84 as from 72. There are 22 measurements at 72 hours and 14 measurements at 84 hours, so we take as our degrees of freedom *min* (22-1,14-1) = 13. We therefore obtain prediction intervals by the same method as before of (14.497,18.297) for *m* = 5 and (14.979,17.815) for *m* = 30.

For completeness, Table 2 provides the sample means and variances (when they exist) for the cases for which we had between one and five measurements (we have not formulated prediction intervals in these cases). Any remaining time, temperature, and instar combinations that do not appear in either the above or below tables are ones for which we have no measurements, and for these the interpolation method described above should be used to form prediction intervals, if necessary.
